# Discovery of novel therapeutic targets in cancer using patient-specific gene regulatory networks

**DOI:** 10.1101/2022.01.31.478503

**Authors:** Andre Neil Forbes, Duo Xu, Sandra Cohen, Priya Pancholi, Ekta Khurana

## Abstract

Most cancer types lack effective targeted therapeutic options and in cancers where first-line targeted therapies are available, treatment resistance is a huge challenge. Recent technological advances enable the use of ATAC-seq and RNA-seq on patient biopsies in a high-throughput manner. Here we present a computational approach that leverages these datasets to identify novel drug targets based on tumor lineage. We constructed patient-specific gene regulatory networks for 371 patients of 22 cancer types using machine learning approaches trained using three-dimensional genomic data for enhancer to promoter contacts. Next, we identify the key transcription factors (TFs) in these networks, which are used to identify therapeutic vulnerabilities either by direct targeting of TFs or proteins that they co-operate with. We validate four novel candidates identified for neuroendocrine, liver and renal cancers, which have a dismal prognosis with current therapeutic options. We present a novel approach to use the increasing amounts of functional genomics data from patient biospecimens for identification of novel drug targets.

## Introduction

Targeted therapy has dramatically improved the survival outcomes of many cancers, for example the use of imatinib in chronic myelogenous leukemia, vemurafenib in advanced melanoma, tamoxifen in breast and cetuximab in metastatic colon cancer^1–4^. However, most cancer types currently lack effective targeted therapeutic options and even the cancers where first-line targeted therapies are available, the prevalence of treatment resistance and inevitable disease progression is a continuous challenge^5–7^. As such, the discovery of novel targeted agents, particularly in aggressive and to date, untreatable diseases, is a high priority. The most commonly used approach to identify drug targets is based upon the presence of genotypic alterations^8,9^.However, not all tumor types and subtypes exhibit targetable genetic alterations. With this in mind, a recent high-throughput approach was undertaken to repurpose existing compounds by screening thousands of compounds in cell lines^10^. While this approach is powerful, it usually results in a large number of potential hits both in compounds and druggable gene targets. In fact, in the most comprehensive drug screening resource available, the average cell line was highly sensitive to over 200 drugs targeting ~250 genes.

It is known that tumors from different cancer types may evolve towards similar lineage driven by the same master regulators or transcription factors (TFs) and indeed lineage plasticity is one of the major causes of resistance to targeted therapies^11–14^. While master regulators can be identified using gene regulatory networks constructed using RNA-seq data and have revealed successful drug candidates such as mirdametinib in breast cancer, they cannot infer directionality of the interactions and do not leverage the chromatin accessibility data that can now be generated for patient samples^15–18^.

Recent advances in technology allow the use of ATAC-seq to identify open chromatin regions directly from patient biospecimens ^19^. Integration of ATAC-seq and RNA-seq data allows construction of directional regulatory networks^20,21^. While previous methods for regulatory network construction modeled the relationship between chromatin accessibility and gene expression in simplistic linear correlation-based methods, recently we have developed a computational method that models the joint contribution of multiple ATAC-seq peaks to gene expression using a random forest model resulting in more accurate patient-specific enhancer to target gene connections^22^. Here, we find that the increased accuracy of enhancer to target gene connections results in more accurate regulatory networks (with nodes consisting of TFs and target genes) and identification of key TFs. While the DNA binding activity of TFs can be hard to target therapeutically, it has been shown that identification of master TFs can reveal other potential targets, i.e. the proteins that co-bind/interact with them^23,24^. This enabled us to develop an approach, CaRNets (Cancer Regulatory Networks and Susceptibilities), that uses knowledge of gene regulatory networks to predict drug targets based upon identification of key TFs and their interacting proteins. Application of CaRNets on 371 patient samples across 22 cancer types revealed 22 clusters, 13 of which consist of multiple cancer types. Importantly, our approach revealed novel drug targets for all 22 clusters. Among these neuroendocrine, hepatocellular, and renal cancers are aggressive cancers with no first line targeted therapies. CaRNets predicted NAMPT and PPAT inhibitors for neuroendocrine cancer, an MDM2 inhibitor for renal cancer and a BRD9 inhibitor for hepatocellular cancer as effective therapeutic options, all of which were validated using cell viability assays showing the strong promise of our approach.

## Results

### Patient clusters based on TF scores

Patient subtyping based on molecular features can be done using RNA-seq or ATAC-seq. We find that while the total number of clusters is similar using unsupervised methods on these data types, often the assignments of patients to the clusters differ (Supplementary Figure 2A). To overcome this, we conceptualized an approach that enables us to identify features that integrate both accessibility and expression datasets as well as the network structures enabled by our peak-gene predictions. We constructed regulatory networks for 371 patients from 22 cancer types using ATAC-seq and RNA-seq data^20^. The enhancer to gene edges were constructed using a random forest model that was trained using three-dimensional CTCF/cohesin ChIA-PET data and models the joint contribution of multiple ATAC-seq peaks to gene expression while accounting for confounders such as copy-number variations^22^. We then identified TF footprints at accessible regions and built edges between TFs and their target genes ^25,26^ (Figure 1A). We identified likely binding events for 1764 sequence specific motifs corresponding to 850 TFs and find that they bind to 54,153 unique regions predicted to target 17,894 genes. We obtain high area under precision-recall curve (AUPRC) using ChIP-Seq data of 20 TFs in MCF7 cells as gold standard demonstrating the accuracy of TF binding predictions (Supplementary Figure 1) ^28,29^. For validation of the gene targets of TFs, the genes that are dysregulated upon siRNA mediated knockdown of ESR1 in T47D are significantly enriched for predicted ESR1 targets using our network construction approach (log odds ratio, LOR= 0.32, p-value 7.69×10^-5^) vs. the targets based on the simple correlation-based approach (LOR=0.02, p-value 0.77)^20,30^.

**Figure 1.**
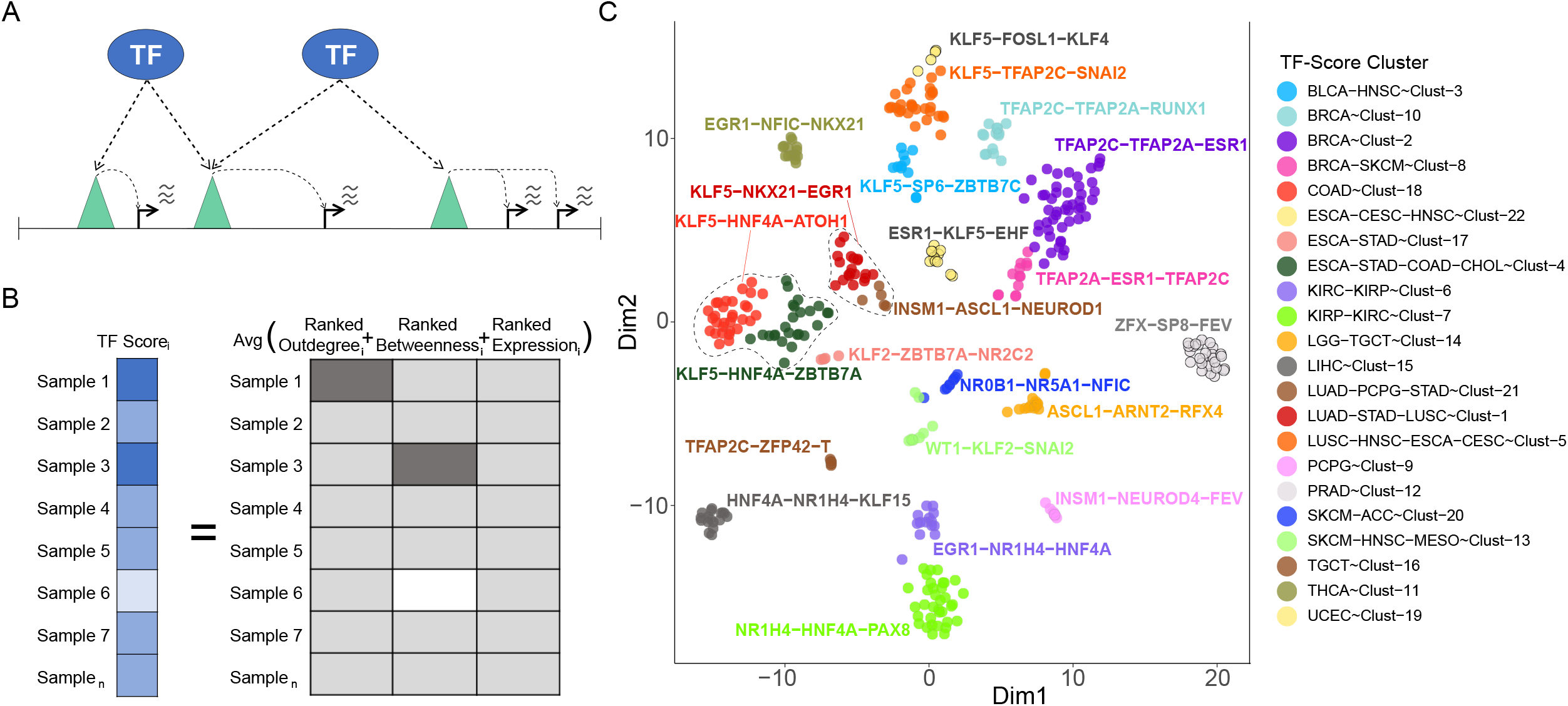
Construction, clustering, and identification of key TFs from patient regulatory networks. A) Construction of edges between TFs and regulatory elements and via those regulatory elements, target genes. B) Schematic of TF-Score for TFs across patients. C) t-SNE of top 15 PCs of TF-scores in 371 patients with 22 identified clusters and their cancer types. Clusters in figure are labeled with the Top 3 TFs in that cluster by relative TF-Score.

Next, we used a combination of three features to generate a score for each TF in each patient: outdegree centrality (number of regulatory targets of the TF), betweenness centrality (frequency of a TF falling on the shortest paths between other TFs and their regulatory targets to identify bottlenecks) and the TF expression based on RNA-seq (Figure 1B) (**Methods**). These TF scores represent the importance of each TF in each patient-specific network and were then used to cluster the patient samples based on an unsupervised, density-based approach (**Methods**)^31–33^. The rationale behind using TF scores to cluster samples is to identify features representative of cell lineage and phenotype while enabling us to identify the key regulators responsible for maintaining that lineage. We identify 22 patient clusters with a median size of 14 patients and a maximum of 49 patients (Figure 1C). While 9 out of 22 clusters correspond to only one cancer type, 13 clusters contain patients from multiple cancer types. For example, the largest cluster (BRCA-Cluster2) corresponds to non-basal breast cancer samples with TFAP2A/B/C, ESR1, FOXA1, AR, SPDEF and MYB as key TFs -- all of which have previous evidence of importance in Estrogen Receptor positive (ER+) breast cancer ^34–37^. Similarly, we identify ESR1, PGR and FOXJ1 as key TFs in Uterine and Endometrial cancer (UCEC-Cluster19) ^38^ and ZFX, FOXA1, HOXB13 and ETS-family members SPDEF and FEV as key TFs in prostate adenocarcinoma (PRAD-Cluster12)^39–43^ (Figure 1C, top 3 TFs in each cluster are shown and Supplementary Fig. 3).

While the patient clusters based on TF scores and discussed above are similar to those based on ATAC-seq and/or RNA-seq data alone, we identify some distinct clusters that stand out only using TF score-based clustering (Supplementary Fig. 2 and Supplementary Table 2). One example is a cluster (LUSC-HNSC-ESCA-CESC-Cluster5) of squamous-like cancers from lung squamous cell carcinoma (LUSC), head and neck squamous cell carcinoma (HNSC), esophageal carcinoma (ESCA) and Cervical squamous cell carcinoma and endocervical adenocarcinoma (CESC) patients, in which we identify TP73, TP63 and SOX2 as key TFs, a finding corroborated by other studies (Supplementary Fig. 3)^12,20,44^. Another example is LUAD-PCPG-STAD-Cluster21 which contains samples that were annotated as lung adenocarcinoma (LUAD), stomach adenocarcinoma (STAD) and pheochromocytoma/paraganglioma (PCPG). Notably, PCPGs are a known neuroendocrine cancer which raised the possibility that this cluster contained other neuroendocrine samples that had not previously been annotated^45^. We identify known neuroendocrine markers such as ASCL1, INSM1, SIM1, NEUROD1 and PROX1 as key TFs in this cluster with ASCL1 and NEUROD1 having been previously identified as key TFs in neuroendocrine prostate cancer and small cell lung cancer (SCLC) (Figure 1C)^46,47^. We also observe relatively high expression of known marker genes of neuroendocrine and/or neural-like differentiation such as *SYP, PCSK1* and *CHGA* relative to all other LUAD, STAD and LUSC samples. Comparison of a set of neuroendocrine marker genes across all cancer types further reinforces this observation, with Cluster 21 showing the highest average expression of these genes (z-score:1.93, p=0.0052) across the 22 clusters identified (Figure 2A). As additional validation, the two other neural/neuroendocrine clusters show similarly elevated expression of these genes, low-grade glioma (LGG-TGCT-Cluster14) (z-score:1.59, p=2.71 × 10^-12^) and Cluster-9 PCPG (z-score:1.53, p=3.08 × 10^-9^)^11,48^. Additional validation for this cluster comes from pathological examination of the STAD member of this cluster with the sample assessed as possessing a *“mixed gastric adenocarcinoma and small-cell neuroendocrine phenotype”*

**Figure 2:**
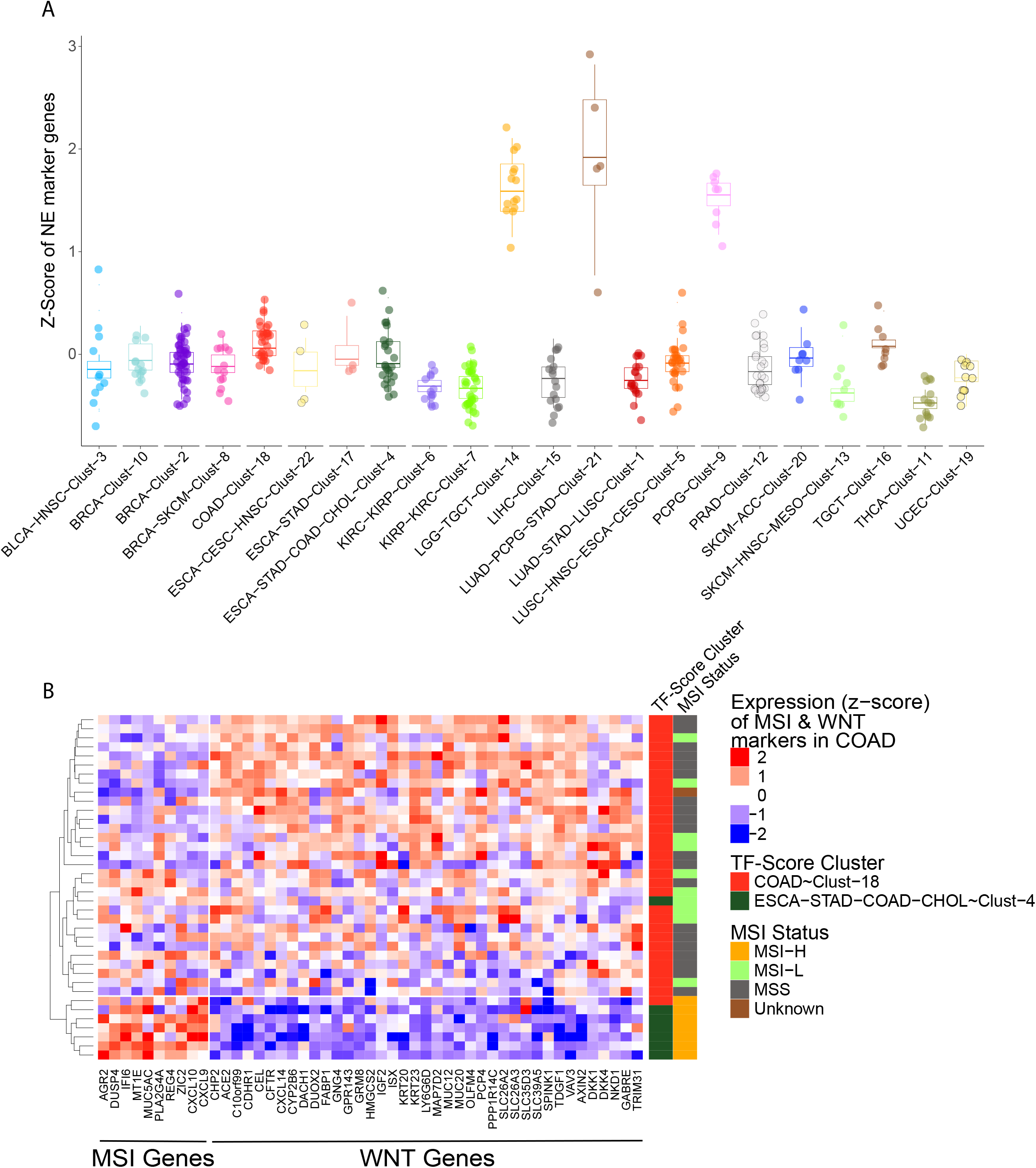
Features of cancer subtypes using TF-Score. A) Expression of neuroendocrine marker genes across 22 clusters identified using TF-Score. Expression of each gene in log2TPM converted to z-scores across all 371 samples. Mean expression of all marker genes plotted on y-axis for each sample. B) Expression of known marker genes for WNT activation and MSI-H cancer phenotypes in colon adenocarcinoma patient samples. WNT activation is commonly seen in COAD cancer.

ESCA-STAD-COAD-CHOL-Cluster4 is yet another distinct multi-disease cluster observed only using TF scores and includes 28% (7/25) colon adenocarcinoma samples (COAD) and the remainder a mixture of cholangiocarcinoma 8% (2/25), esophageal carcinoma (ESCA) 20% (5/25) and stomach adenocarcinoma 44% (11/25). Comparison to Cluster18, which consists of exclusively COAD samples showed that the COAD samples in Cluster4 featured 85% (6/7) of the Microsatellite Instability High (MSI-H) tumors in the entire COAD cohort and were primarily sampled from the proximal colon, observations not mirrored in the unsupervised ATAC-Seq or RNA-Seq clustering in isolation^49^ (Supplementary Figure 2B and C). Consistent with this, we observe higher expression of MSI-associated genes and lower expression of WNT activation markers indicating a convergence of regulation of those genes in the TF space (Figure 2B)^50,51^. Additionally, we see enrichment of gene sets associated with MSI-H in the COAD samples in this mixed cluster relative to other COAD samples (Supplementary Figure 6)^52,53^.

Another multi-disease cluster observed exclusively using TF scores consists of a mixture of ESCA and STAD samples (ESCA-STAD-Cluster17) and is separate from the previously discussed GI cluster, all of which show low somatic copy number alterations in contrast with the ESCA and STAD samples in the cluster (ESCA-STAD-COAD-CHOL-Cluster4) containing the MSI-H colon samples^54^.

Our results show that TF score-based clustering can capture integrated features of the chromatin and transcriptomic landscape and helps in identification of key TFs and master regulators of each cluster. It allows grouping of patients based on similarity of the key regulators, providing an ideal framework to identify potential novel drug targets.

### Cancer Regulatory Networks and Susceptibilities (CaRNetS): Computational framework to identify drug candidates for each cluster

We used the clusters based on TF scores to identify new potential drug targets in a novel approach called Cancer Regulatory Networks and Susceptibilities (CaRNetS). Recent studies have shown that besides the key TFs, the proteins that interact with them can be promising drug targets due to their ability to dysregulate the downstream TF targets, for example the FOXM1:CENPF regulon in prostate cancer, TWIST1/SNAI1:TGFβ and RUNX1/ETS1:CBFβ in metastasis in multiple cancers ^18,55–57^. Thus, to identify the proteins that co-operate with the key TFs in the appropriate biological context, we used the co-dependency scores of genes from DepMap^58^. To do this we first need to identify cell lines that best model patient clusters and identify the targetable genes that are highly correlated with cluster-specific key TFs. We implemented an approach modeled on the nearest-template prediction algorithm using RNA-Seq data from both the patient clusters and cell lines from the Cancer Cell Line Encyclopedia (CCLE) (Figure 3A) (**Methods**)^59^. We applied this approach to 496 cell lines belonging to similar tissue types as the cancer types in our patient cohort; 335 of which we can robustly assign to a patient cluster (Figure 3B).

**Figure 3:**
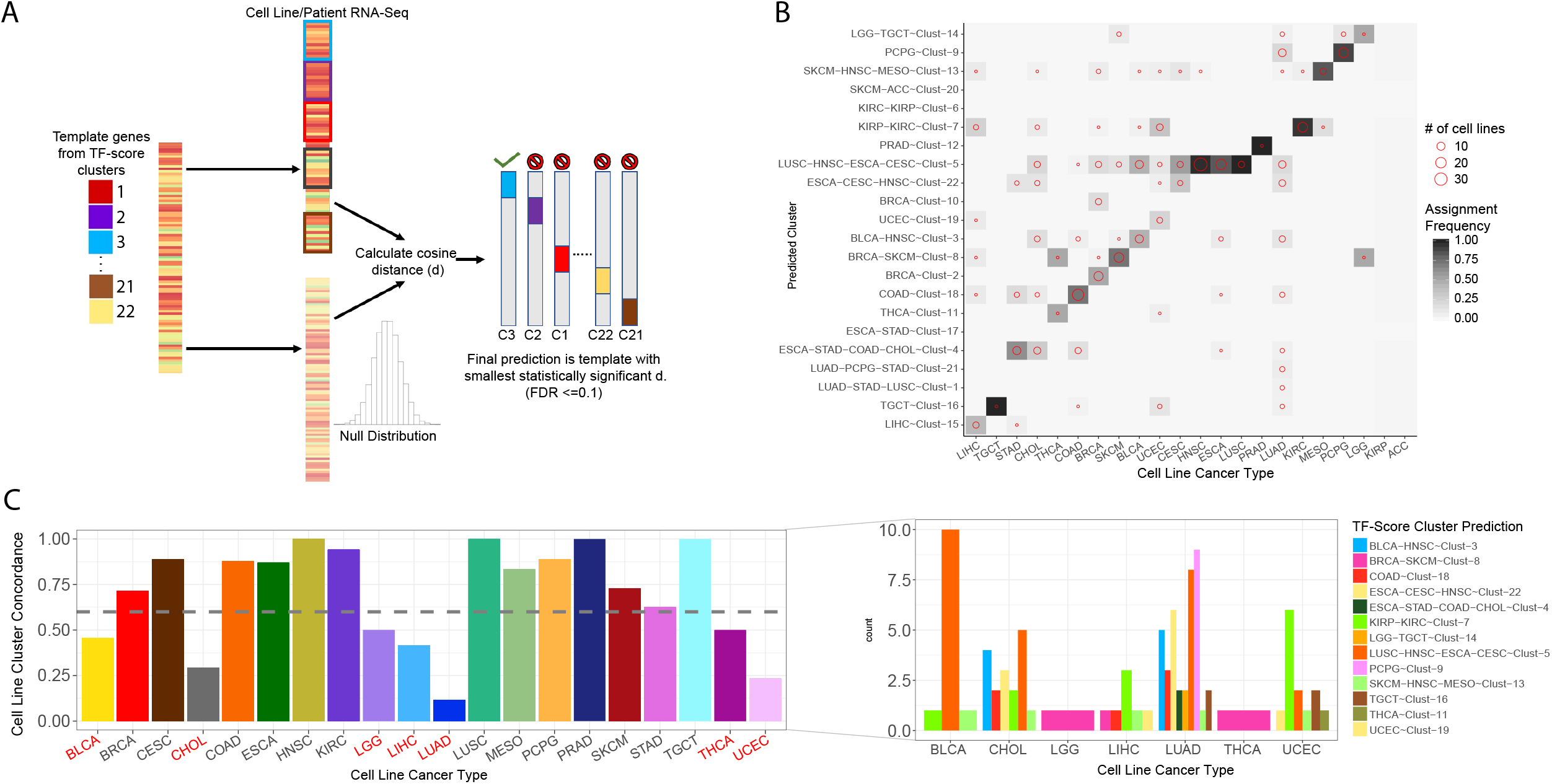
A) Assignment of cell lines to representative clusters using a nearest template prediction based approach.) B) Detailed assignments of 335 cell lines present in both DepMap and CCLE datasets to TF-Score clusters by cell line cancer type. Heatmap is normalized by total number of cell lines in that cancer type. C) Concordance of NTP approach to assigned cell lines to patient clusters by cancer type. Detailed assignments of cancer types with less than 50% concordance showing likely phenotypic convergence of LUAD cell lines to squamous-like and neuroendocrine cancers. Uterine cancers also show significant similarity to renal cancers likely arising from their shared lineage as mesoderm-derived cell types.

The majority of cell lines (64%) are assigned to clusters containing their specific cancer type though there is significant variation across cancer types with a median of 78% cell lines of a given cancer type showing concordance with the cancer type of its assigned cluster (Figure 3C). Prominent examples of divergence from expected cancer types include the cell lines FU97, NCI-H226, SW-1710 and JHH1, which have previously been associated with STAD, LUSC, BLCA and LIHC assigned as LIHC, MESO, KIRC/KIRP and UCEC respectively, an observation supported by other comprehensive approaches^60^. The cancer type with the most divergent assignments from expectation is LUAD. However, we find 37% (32/86) LUAD cell lines predicted to represent non-LUAD clusters are assigned to clusters containing squamous-like cancers and an additional 38% (33/86) of these “misclassified” LUAD cell lines are predicted to represent neural/neuroendocrine-like cancer types, with 94% of those (31/33) having been annotated as small-cell/neuroendocrine-like. This indicates that this RNA-seq based assignment of cell lines may reflect the underlying biology better than the original cancer type assigned to the cell line.

The assignment of cell lines to patient clusters allowed us to use the data from DepMap to identify proteins that likely co-operate with the key TFs in each cluster^24^. To identify genes highly correlated with our key TFs in a cluster-specific fashion that are druggable by existing compounds, we computed correlation of gene dependency scores in a cluster-specific fashion. Next, to identify therapeutically relevant genes, we utilized the Drug Interaction Database to identify those genes with known interactions with an available and/or published compound^61^. We further prioritized TF-Gene pairs using an estimate of drug toxicity of compounds that interact with the gene member of the pair and TF-score from our regulatory networks (Figure 4A) (**Methods**). We integrate these features to create a CaRNetS score for each TF-Gene pair which was used to identify potential drug targets.

**Figure 4:**
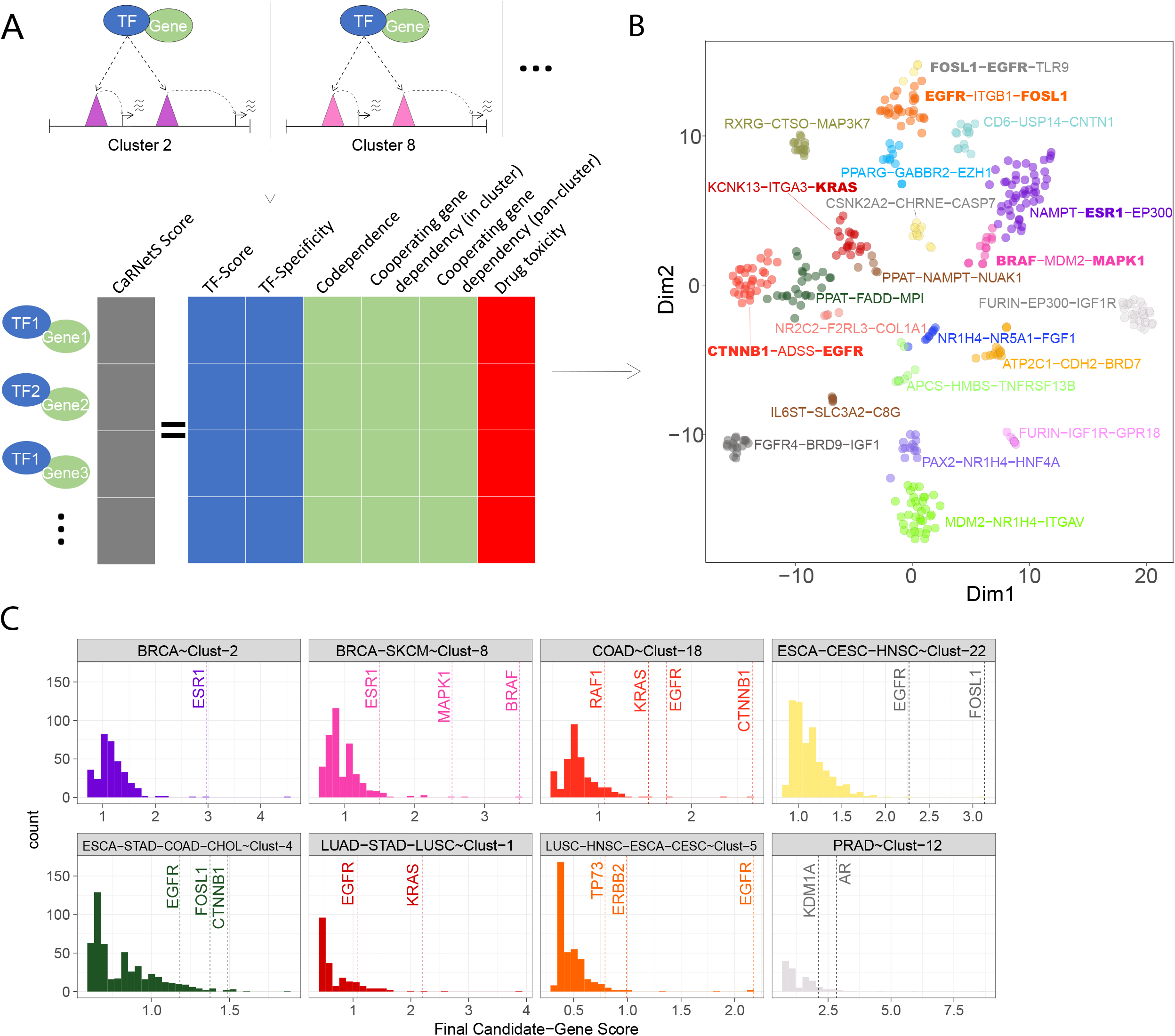
CaRNetS workflow. A) CaRNetS scoring schema showing the features used in druggable gene prioritization. B) Top candidates for each cluster with known candidates highlighted. C) Known candidate genes in 8 clusters and associated CaRNetS scores.

### Drug targets identified using CaRNetS

Our approach identified a set of target genes and corresponding drugs as candidates for each cluster (Figure 4B, Supplementary Table 3). Some of these candidates are known to be drivers of cancers or targets of inhibitors either currently in clinical trials or already approved as top candidates providing validation to the approach (Supplementary Table 5,^2–4,14,62–65^). Indeed, we find that 90% of gene candidates targeted by drugs that are currently used in the clinic or are under trials in their respective cancers are ranked in the 95^th^ percentile or higher of candidates predicted and frequently in the top 20 candidates by absolute rank (Figure 4C). For example, ESR1 in ER+ breast cancer, BRAF in melanoma, B-Catenin in colon and other GI adenocarcinomas, EGFR in both squamous and non-squamous lung cancer and AR in prostate cancer (Figure 4b-c). Importantly our approach reveals other potential targetable genes for each cluster, which is especially promising for those with few or no targeted therapies such as neuroendocrine cancers, Testicular and germ cell tumors (TGCT) and mesothelioma (MESO) (Figure 4b). Even for cancers with known targets, resistance to targeted therapy is widespread and thus the novel candidates identified by CaRNetS may offer promising opportunities for combination and/or second-line therapies^5,6^.

We identified EP300 as a top candidate in ER+ BRCA and in PRAD. EP300 is itself a histone acetyltransferase and is frequently found at active enhancer regions. Our result is supported by a recent study that demonstrated significant benefits of EP300/CBP inhibitors in AR--positive BRCA and AR-positive PRAD cell lines with negligible effects on IC50’s for AR--negative cancers^66^. We also identified FGFR4 in LIHC with recent studies showing strong evidence to suggest the efficacy of an inhibitor, Roblitinib, in vitro and in vivo with xenograft models providing strong support to our results^67^. We find other chromatin modulators such as BRD7 in low grade glioma (LGG), BRD9 in Liver/Hepatocellular carcinoma (LIHC) and KDM5A in TGCT, KDM1A, KDM4A, KMT5B and KDM5C in PRAD as high priority druggable genes.

In addition to the above candidates with some evidence from other studies supporting their prioritization, we identified several novel candidate genes in cancers with few effective targeted therapies, namely BRD9 in LIHC, MDM2 in renal cancer (KIRC), and NAMPT and PPAT in neuroendocrine cancers. To validate these predictions, we tested inhibitors of BRD9, MDM2, NAMPT and PPAT in HepG2, A498 and DMS53 cell lines respectively. We observed statistically significant effects on cell growth for BRD9, MDM2, NAMPT and PPAT inhibitors as single agents compared to control cell lines as measured by cell viability assays (Figure 5 a to d) and an inability to identify IC50s for these inhibitors in control cell lines. Further, the observed effect of the NAMPT inhibitor, Teglarinad Chloride, on DMS53 cell lines is concordant with its effect in GOT1 cells, a gut neuroendocrine cell line, both in vitro as well as in xenograft models^68^.

**Figure 5:**
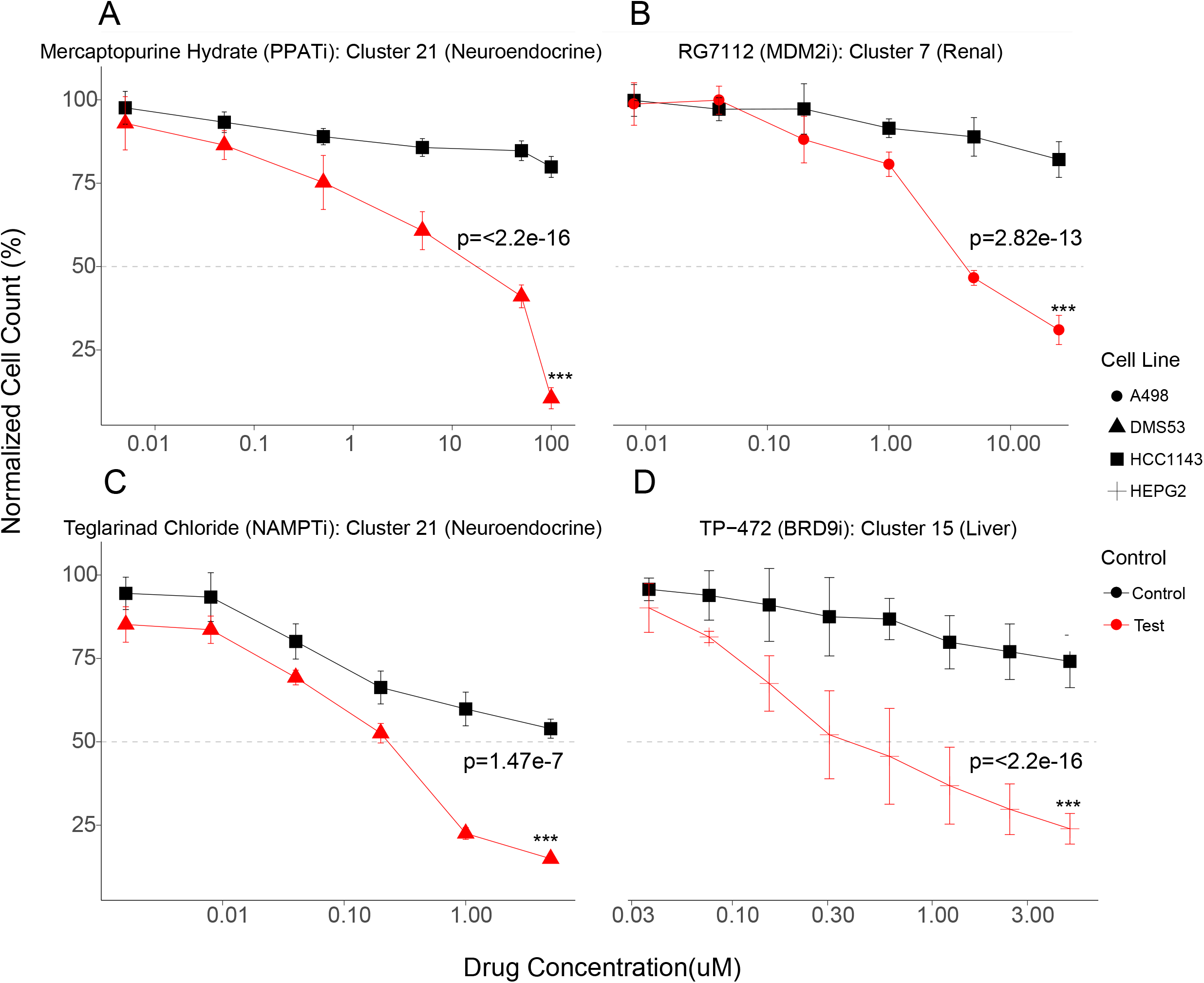
Effect of inhibitors of 4 candidate genes across 3 clusters A) Cell viability assay for Mercaptopurine in DMS53 cells. B) RG7112 in A498 cells. C) Teglarinad Chloride in DMS53 cells. D) TP-472 in HEPG2 cells.

Mercaptopurine hydrate is identified as one of the two drugs targeting the protein PPAT and is highly correlated with both INSM1 and ASCL1, key TFs in neuroendocrine/small cell tumors. The drug has a primary indication in hematological malignancies with occasional use in Crohn’s disease^69–71^. This is consistent with the observation of Balanis et al where they noted some shared susceptibilities between hematological malignancies and neuroendocrine/small-cell tumors^11^. RG7112 is a highly selective inhibitor of MDM2, an oncogenic ubiquitin ligase, responsible for ubiquitination and degradation of TP53 and other tumor suppressor proteins^72–74^. MDM2 is highly correlated with PAX2 in renal cell lines, a gene implicated in other renal pathologies^75,76^. Our networks indicate that PAX2 is regulated by NR2F2, another key TF in renal cancers. Teglarinad Chloride was identified as the only drug targeted at Nicotinamide phosphoribosyltransferase (NAMPT) a key member of the NAD+ salvage pathway with NAMPT, a direct regulatory target of ASCL1 itself in our regulatory networks ^77,78^. TP-472 was one of several highly selective inhibitors of BRD9, identified as a top drug candidate in hepatocellular cancer. BRD9 is a bromodomain protein and has been characterized as part of non-canonical BAF (SWI/SNF) complexes that function as ATP-dependent chromatin remodelers^79^. In our analysis it is highly correlated with HNF4A, which has the highest TF-score in hepatocellular cancers and importantly, BRD9 is both highly expressed in LIHC and associated with worse outcomes^80^.

Importantly, all of the four drugs validated here are candidates in cancer types that have dismal outcomes even with the use of existing treatment options (Figure 5). In LIHC, the patients with the best outcomes can expect a 5-year survival of 50-70% with a recurrence rate of up to 80% at 5 years depending on initial treatment modality^81–83^. Similarly, neuroendocrine lung cancers have dismal outcomes with ~32% 5-year survival rates^84–86^. This underscores the dire need for effective first line therapies for these cancers and the utility of CaRNetS towards this goal.

## Discussion

We present a novel approach that leverages a high-quality set of enhancer-gene predictions in a large patient cohort to reconstruct directed TF-Gene networks for 371 primary cancer patients across 22 cancer types. We compute scores for all TFs that reflect their importance at the integrated levels of chromatin accessibility and transcription and allow clustering of patient samples. While some of these TFs can be potentially targeted by drugs, in general, the DNA binding activity of TFs is hard to target. Thus, we develop a framework which relies on the interactions of TFs with other targetable proteins in the appropriate biological context. Importantly, our approach, CaRNetS, identifies known key genes used for targeted therapy, such as KRAS and EGFR in lung and colon adenocarcinomas, ESR1 in ER+ breast cancer, AR in prostate cancer and multiple others. With these known positive cases as a reference point, we sought to validate the top candidates in cancers with limited effective targeted therapies. Validation was performed on 4 gene candidates across 3 disease types. Our approach found that inhibition of all 4 candidate genes significantly affected cell growth in cell lines representing renal, liver, and neuroendocrine cancer types as compared to controls.

We note that although the essentiality scores of genes in these cell lines also indicate molecular dependencies and the sole use of cell line dependency data can also help identify drug candidates, a major limitation is the volume of genes identified as essential for a given cell line with the average cell line dependent on the presence of ~2150 genes. High throughput screens are an incredible tool that enable broader testing and identification of key pathways previously unexplored in many cancers. However, there exists the potential for false negatives due to issues with dosing amounts and scheduling (chronic treatment vs single treatments) while seemingly positive drugs may be highly toxic in vivo. We intentionally consider drug toxicity as an important feature in CaRNetS, a feature typically ignored by most gene dependency-based approaches. CaRNetS uses an approach that relies on the molecular features of tumors and may be used in integration with CRISPR-based approaches to identify the most promising candidates. The TFs identified represent key contributors to the cancer type/cell lineage and co-dependent genes are typically members of the same signaling cascades and pathways, are protein-protein interactors, and/or regulatory targets. CaRNets is available online at GitHub and a docker image has been made available for download.

We envision that with the ease of performing ATAC-seq assays on patient biopsies, tremendous amounts of ATAC-seq data will be generated from cancer samples in the future, including for metastatic cancers. CaRNets will enable the use of these datasets to answer the most pressing questions in cancer biology and identify novel drug targets.

**Table 1:**
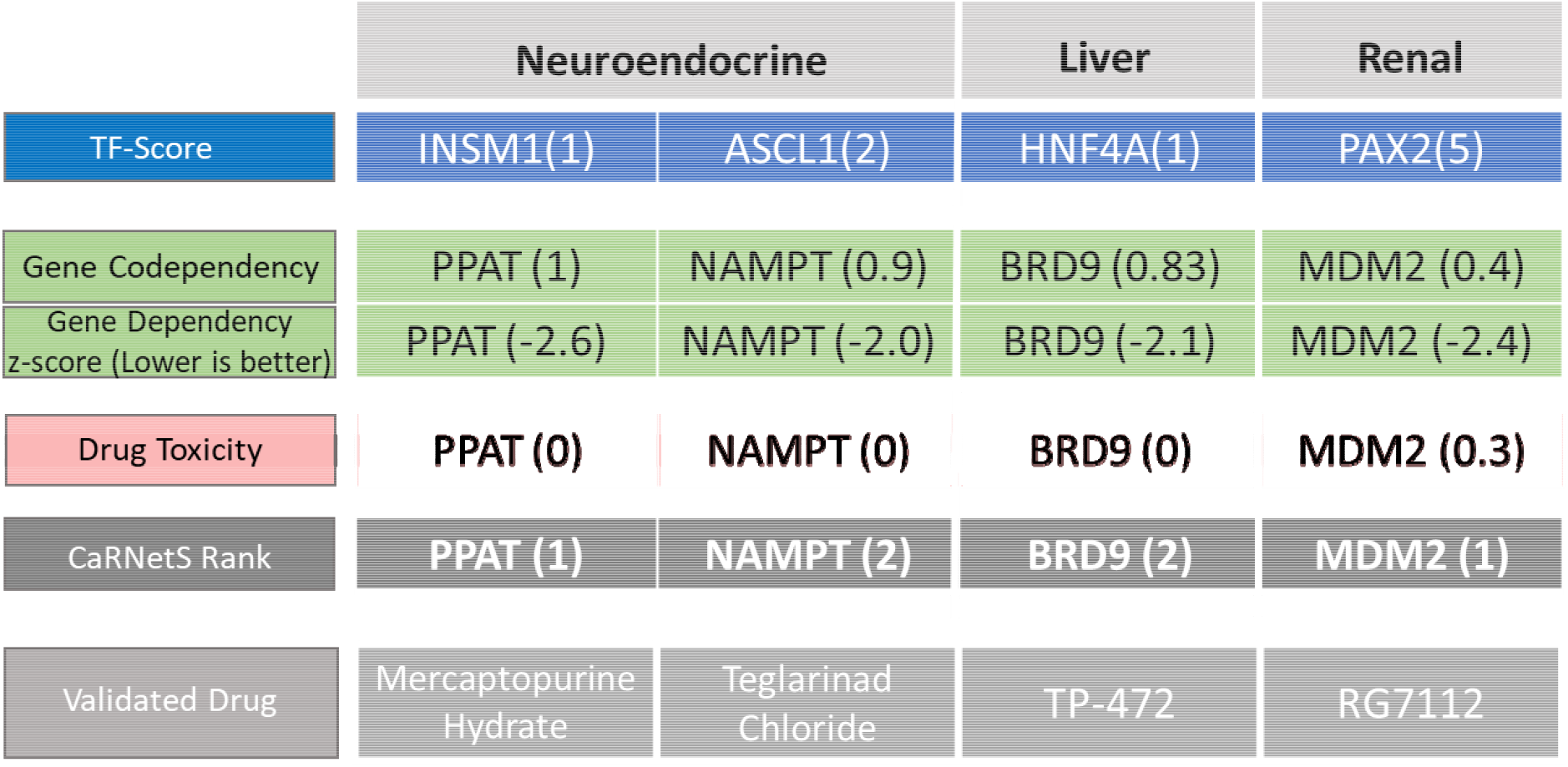
Validated Drugs and their target genes and CaRNets Rank

## Supporting information

Resources Table

---

**Supplementary Figure 1:**
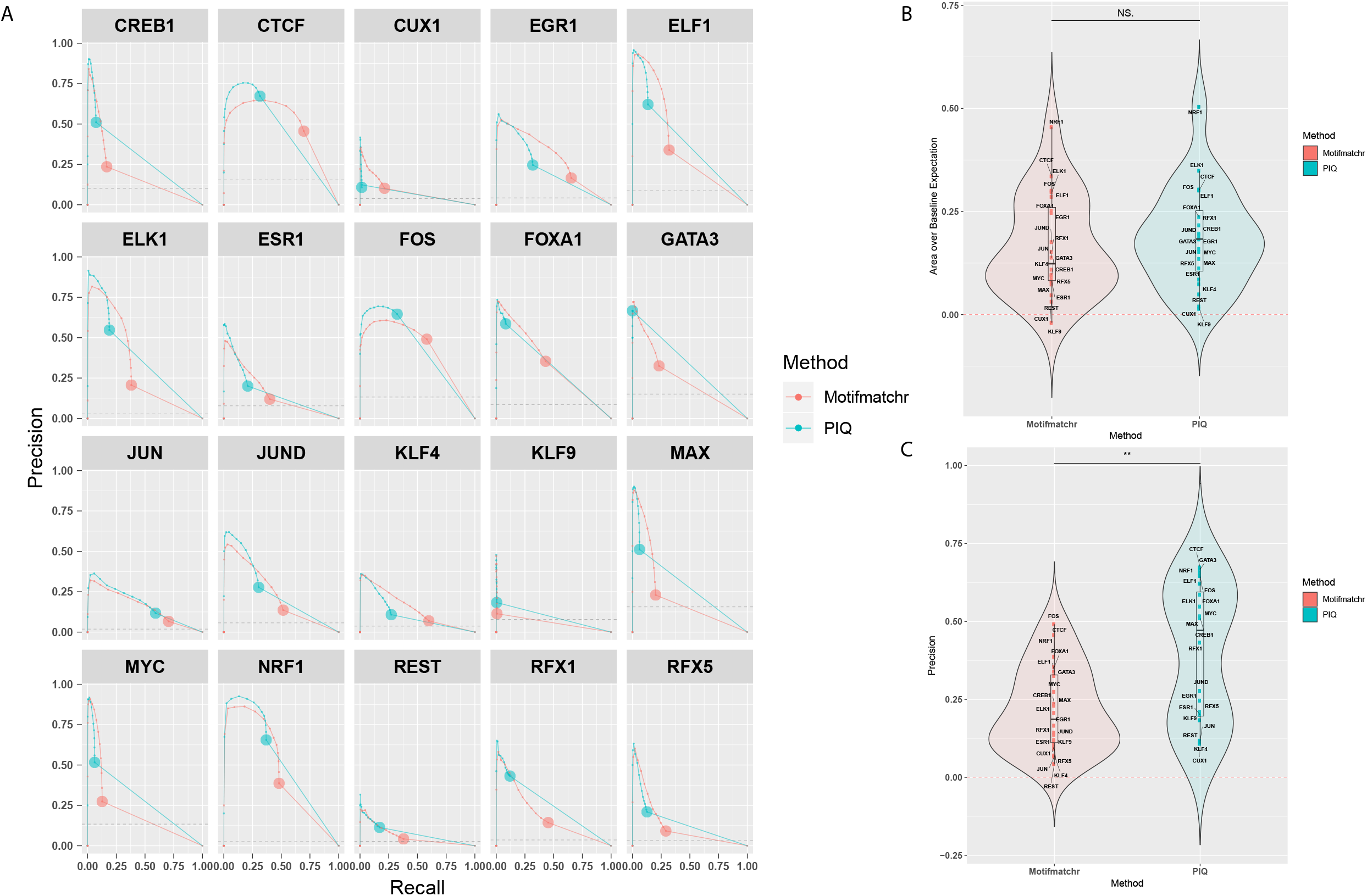
Benchmarking of TF footprinting approach vs ChIP-Seq for detection of true TF binding events in open chromatin. A) Precision-Recall curves for 20 TFs in MCF7 cells at varying peak heights. B) Area under PR-curves above baseline expectation. C) Precision at regions defined as open chromatin for final networks.

**Supplementary Figure 2:**
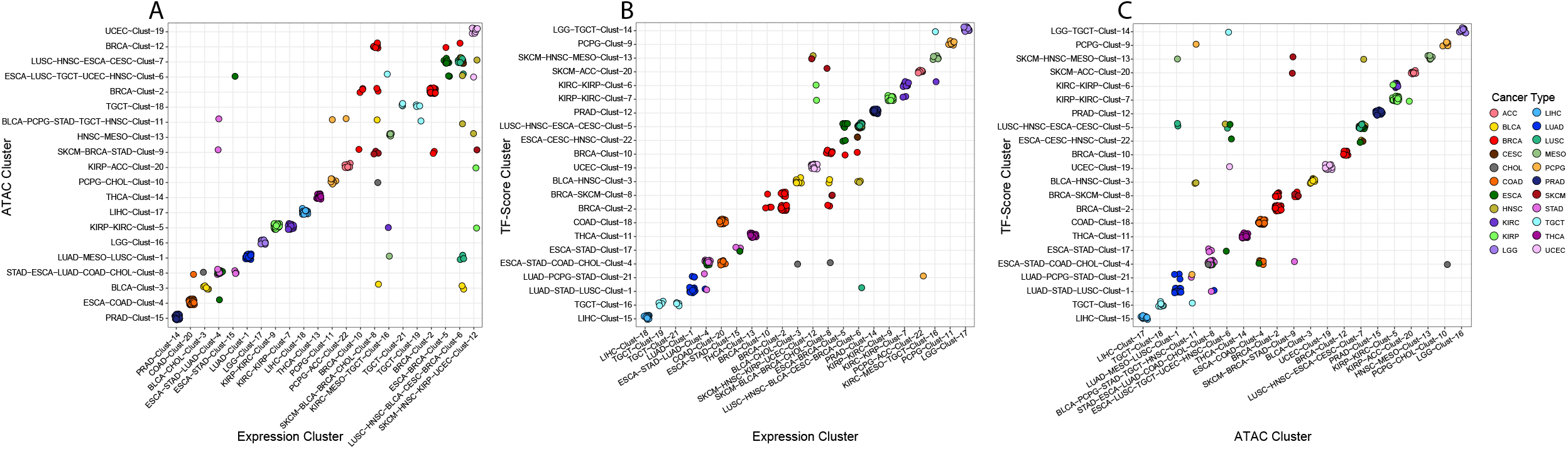
Comparison of TF-score, ATAC-Seq and RNA-Seq based clustering approaches a) ATAC-Seq to RNA-Seq b) TF-Score to RNA-Seq c) TF-Score to ATAC-Seq. Higher concordance observed between TF-Score and ATAC-Seq/RNA-Seq in isolation.

**Supplementary Figure 3:**
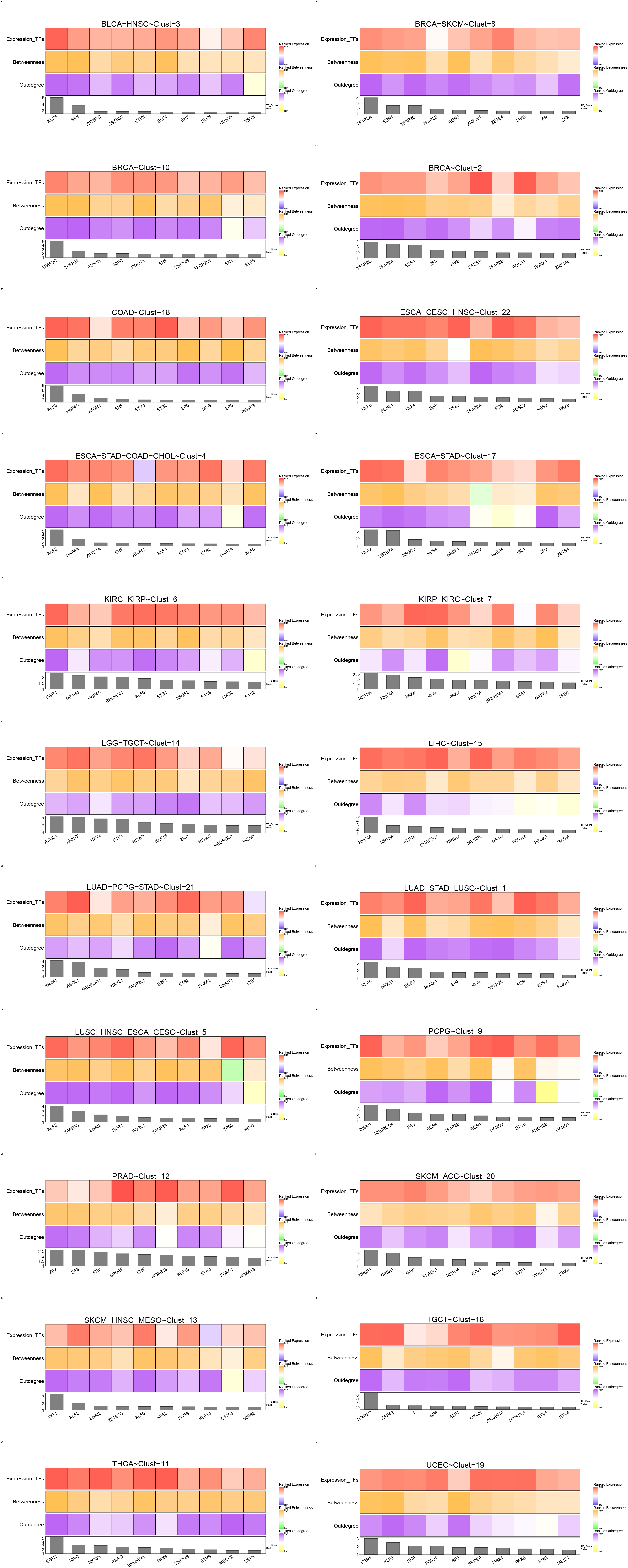
Heatmap of key TFs identified for 22 clusters in TF-score space with the contributions of expression, betweenness and outdegree centrality identified.

**Supplementary Figure 4:**
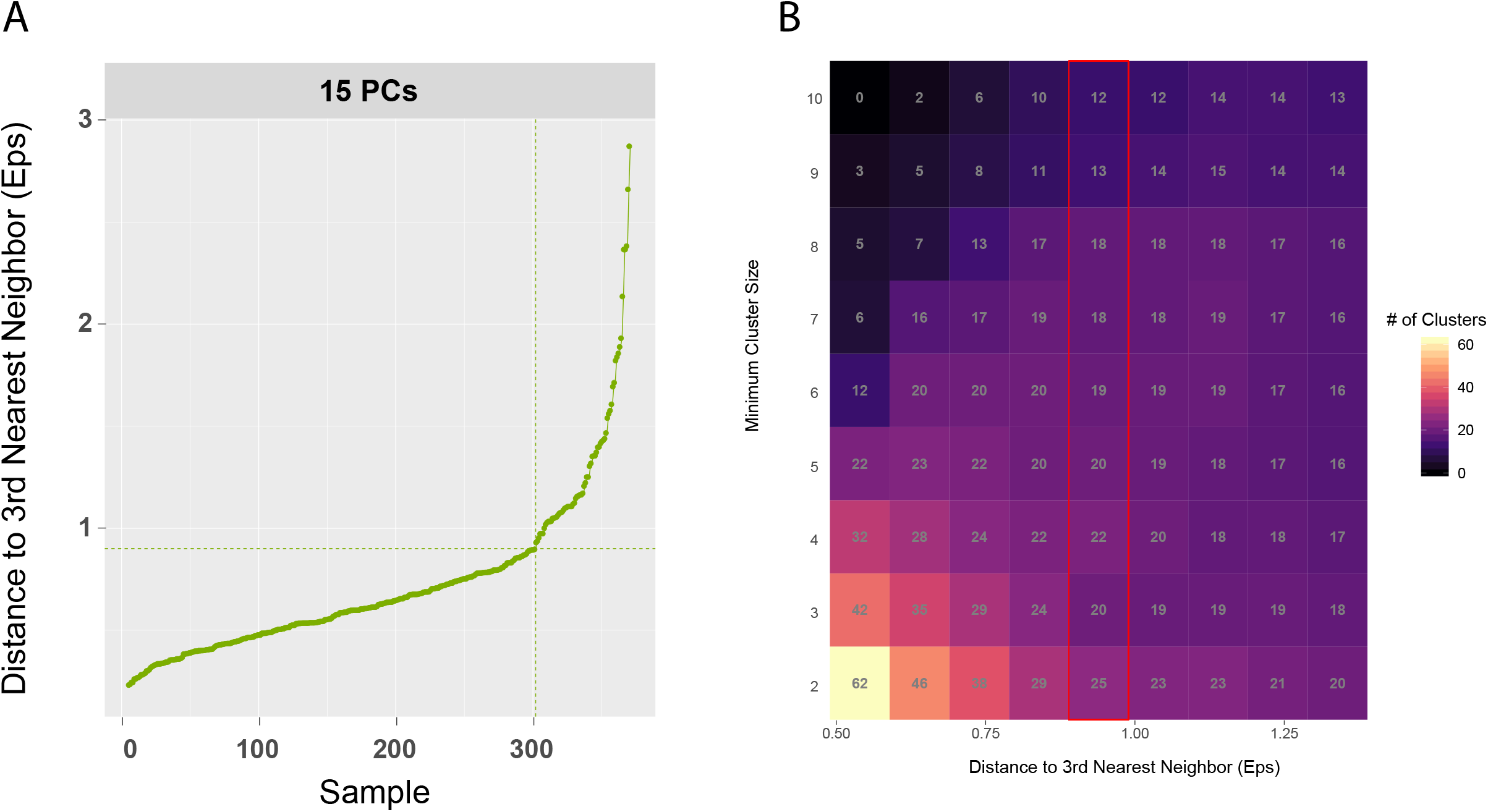
Parameter estimation and tuning for density-based clustering. A) Optimal distance estimation. B) Number of clusters identified at 9 values of eps (+/- 40% of optimal) and 9 minimum cluster sizes.

**Supplementary Figure 5:**
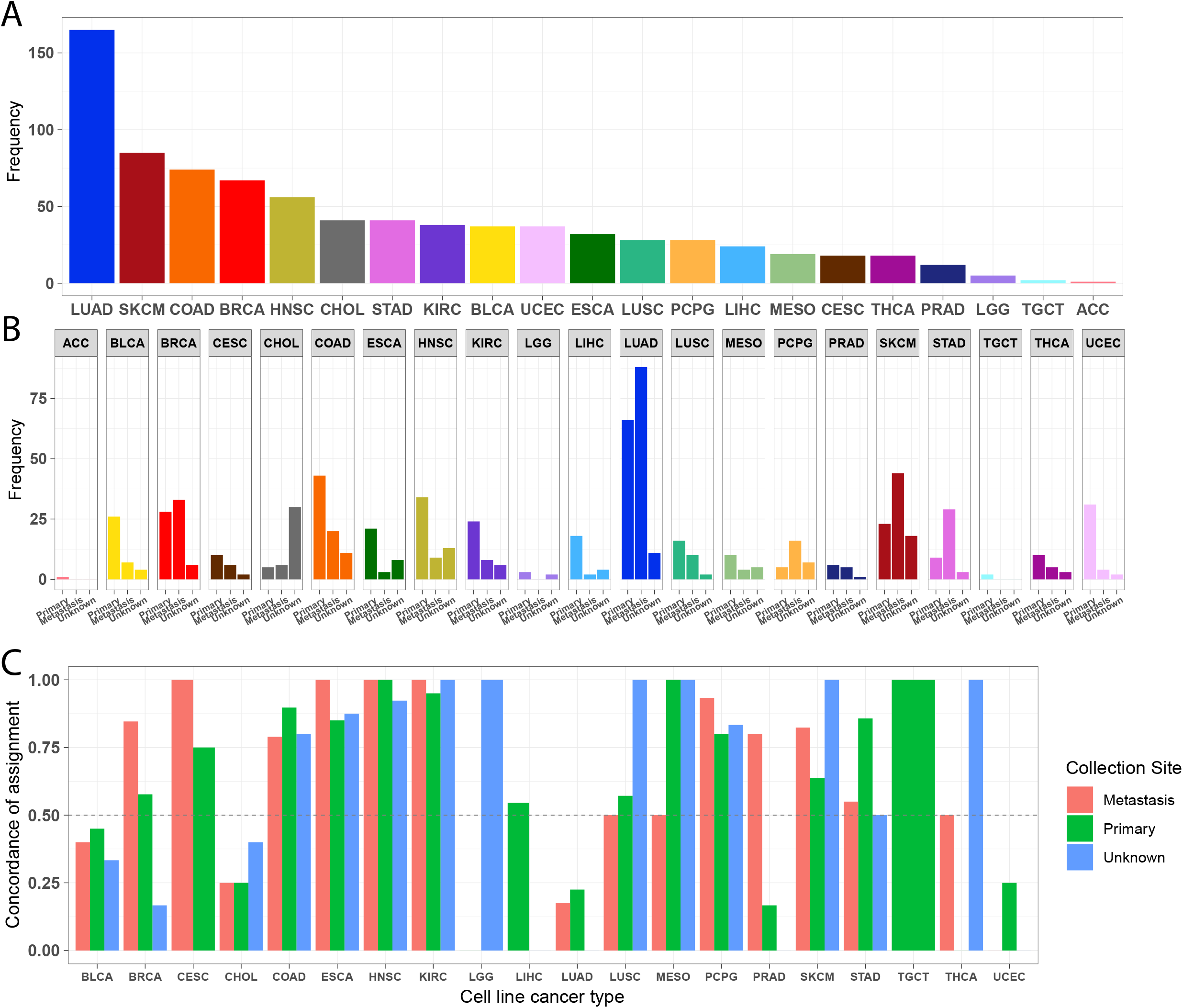
Concordance of NTP-based approach for identifying representative cell line models of patient clusters broken down by sampling site. A) Distribution of cell lines by cancer/tissue type of origin. B)Distribution of cell lines by cancer type and sampling location. C) Concordance of patient cluster predictions with annotated cancer type of cell lines by sampling site..

**Supplementary Figure 6:**
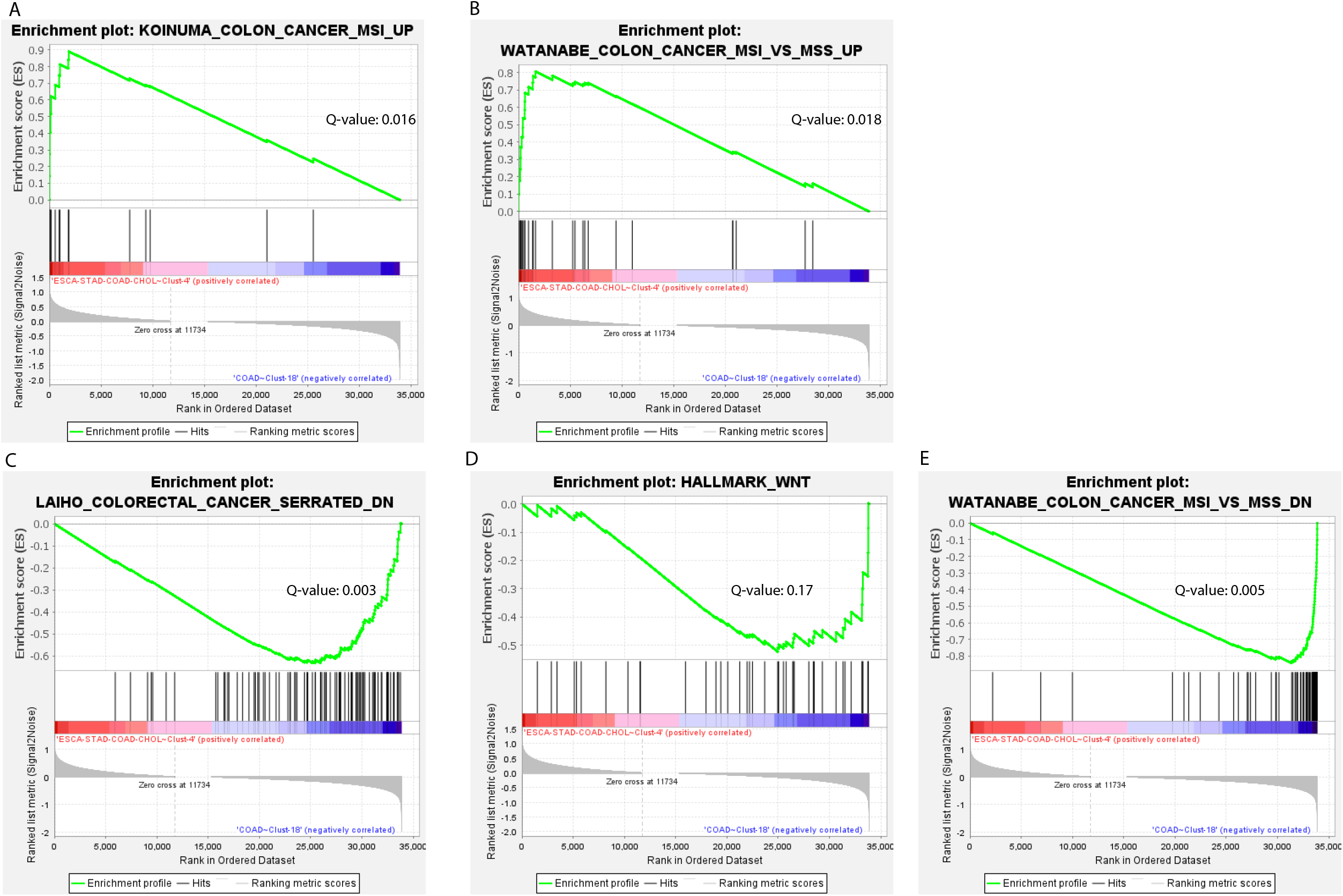
Gene-Set enrichment analysis results for 5 genesets associated with canonical colon adenocarcinoma development and MSI-H status with FDR <=0.2.

Supplementary Table 1: TF-Score cluster predictions for 828 cell lines

Supplementary Table 2 Patient cluster assignments using unsupervised ATAC-Seq, RNA-Seq and TF-Score with associated cancer-specific metadata for all samples.

Supplementary Table 3: Carnets score predictions for top-20 candidates in all 22 clusters

Supplementary Table 4: Encode ChIP-Seq data for 20 TFs used for benchmarking the TF-footprinting approach.

Supplementary Table 5: Genes currently targeted by agents currently in use in the clinic and/or clinical trials by cancer type.

## Methods

### Data processing

ATAC-Seq of 400 primary cancer patients was from Corces et al. ^15^ and RNA-seq data were extracted from NCI Genomic Data Commons (GDC) ^91^ with matching sample submitter_id. The ATAC-seq counts and RNA-seq counts used in this study were preprocessed by the above studies; Log2 of Transcripts Per Million (TPM) was adopted for RNA-seq counts, while log2 counts are used for ATAC-seq counts.

### Cell Culture

MCF7 (HTB-22), MDA-MB-231 (HTB-26) and T-47D cell lines cell lines were obtained from ATCC. MCF7 and MDA-MB-231 were cultured in Dulbecco’s Modified Eagle Medium (DMEM); T-47D was cultured in RPMI Medium 1640.All media were obtained from Gibco and were supplemented with 10% Fetal Bovine Serum (Omega Scientific), 1% Sodium Pyruvate (Gibco), 1% Pen Strep (Gibco), 1% HEPES Buffer (Corning). T-47D media was additionally supplemented with 0.2 units/mL Bovine Insulin (Sigma Aldrich, IO516). Cells were regularly passaged and tested for presence of mycoplasma contamination (Mycoplasma PCR Detection Kit, Applied Biological Materials).

### RNA-seq and ATAC-seq data processing

RNA-seq and ATAC-seq were generated for 3 cell lines (MCF-7, T-47D and MDA-MB-231). Cells were grown to near confluence, pelleted and frozen at −80°C.

RNA was isolated using the RNeasy Plus Mini kit (74134, Qiagen) and sent to the Weill Cornell Medicine Genomics Core facility, where an Illumina TruSeq stranded mRNA Library prep was made using HiSeq 2500 and sequenced on the Illumina NovaSeq 6000 S1 Flow Cell, PE100 cycles. RNA-seq data were aligned to hg38 using STAR, counts was calculated using HTSeq and TPM was calculated as described by Wagner et al.

For ATACseq, nuclei were isolated from >95% viable cells in log phase growth and processed at the Weill Cornell Medicine Epigenomics Core using the OMIN-ATAC protocol as detailed in and sequenced on Illumina NovaSeq 6000, SP Flow Cell PE50 cycles. ATAC-seq data was mapped to hg38 and processed using ENCODE ATAC-seq pipeline (https://www.encodeproject.org/atac-seq/).

### Connecting TF-peak edges

For each patient, we connected the TF to peak links by using a footprint-based method, PIQ ^29^.

Likely TF binding events are identified for 1764 motifs corresponding to 850 TFs and TF families on a per sample basis. In cases where a TF possesses multiple motifs, we use the union of those motifs as the set of binding events for that TF. Binding events are then subset to those with high (>=0.7) estimated positive predictive values (PPV). This final set of binding events is considered as the true set and intersected with known regulatory-elements and their target genes to create an edge list of TFs-peaks-genes for each patient.

### Identifying targets of regulatory elements (DGTAC)

Elasticnet regression is performed for all peaks within the +/-0.5 Mbp distance of the TSS of every gene. We used ElasticNet regression to calculate the coefficient of all peaks within region to the TSS and collected coefficients as peaks’ weight to the target gene expression. After filtering out peaks with zero coefficient, for each peak of each gene, we calculate a residual I term representing the contribution of the single peak to the target gene expression. We construct a feature matrix consisting of peak heights, peak residuI(*e*), distance to TSS, copy number alteration of presumed gene target and gene expression and train a random forest model to predict gene expression using the aforementioned features. We use ChIA-PET enriched for CTCF/cohesin loops in MCF7 cells as true interacting pairs. The predicted peak-gene links represent the targets of regulatory elements and are consistent across cell lines and cancer types used as gold standards.^22^ Model was applied to T-47D, MCF7 and MDA-MB-231 cell lines to predict peak-gene connections.

### Validating TF-peak edges

To validate TF-Peak edges, TF footprints are intersected with the patient-specific peak-to-gene links for MCF7 then compared to ChIP-Seq binding events for the same TFs in MCF7 within the same regulatory regions. Over the range of lower accessibility peak to higher accessibility peaks, we calculate precision and recall for each TF. True positives (TP) are defined as peaks predicted to be bound by both ChIP-Seq and PIQ. False positives (FP) are defined as peaks predicted to be bound by PIQ but not ChIP-Seq. False negatives (FN) are defined as peaks predicted to be bound by ChIP-Seq but not PIQ. These predictions are compared to the predicted TF binding sites using a motif search performed using motifmatchr in the same regions and for the same range of accessibility^87^.

### Validating Regulatory targets of TFs

RNA-Seq data from T47D siRNA mediated knockdown of ESR1 was collected and log2 fold-change (log2FC) was calculated relative to a non-targeting control. True targets of ESR1 are genes with a log2FC <=-0.5 after siESR1 K/D^30^. We compare DGTAC to a correlation-based method derived from the same patient dataset. To identify TF-binding events we use a footprinting approach to find ESR1 binding events in all open chromatin in an ER+ BRCA cell line (MCF7)^26^. We subsequently intersect these binding events with predicted peak-gene connections for DGTAC and the correlation-based method separately to identify the predicted target genes of ESR1. We use a fisher’s exact test to quantify the magnitude and statistical significance of correctly identifying ESR1 target genes in the siRNA experiment using predicted TF binding events and regulatory targets of enhancer regions. Additional validation of DGTAC can be found in other references. ^22^

### Identifying critical TFs for patient network clusters

Edgelists for each patient consisting of TF-gene edges are converted to graphs and each node evaluated for outdegree and betweenness centrality. Betweenness and outdegree are separately converted to within-sample ranks with the highest value for each metric being ranked 1st. We calculate TF-score as the within-sample, per TF, average of the rank of betweenness centrality, outdegree and gene expression.

*For TFi in sample_x_*

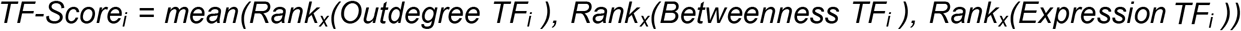

Samples are projected onto a 2-dimensional space using an R implementation of t-distributed stochastic neighbor embedding (tSNE) on the top 15 principal components of the TF-score matrix of all TFs for 371 samples with a perplexity of 30 being used for tSNE. We subsequently cluster samples using an implementation of Density-based spatial clustering of applications with noise (DBSCAN) in R^38,39^. We specify a minimum cluster size of 4 samples and an epsilon that corresponds to the largest changes in the slope of the K-distance curve. Curves used for epsilon selection can be found in Supplementary Figure 4.

Critical TFs for patient clusters were identified using in-cluster vs. out-of-cluster comparisons of TF-score for every TF using a Wilcoxon rank sum test. Candidates are ordered by the ratio of the means of the two populations and filtered for those candidate TFs meeting statistical significance with p-values adjusted for FDR using the Benjamini-Hochberg method.

### Assigning cell lines to patient clusters

Cell lines are assigned to patient clusters by using a nearest template prediction (NTP) approach using CCLE expression data for the cell lines and expression data from the patient. We define a set of template genes for each cluster as the set of genes that are most highly expressed in a given cluster when compared to all other patient clusters. We use this template to calculate the cosine distance (1-cosine similarity) of a given cell line to the template geneset’s expression in each cluster. We compare this distance to the distances from n=100-1000 randomly sampled genesets of the same length as each template to obtain p-values associated with each predicted distance. Cell lines are then assigned to the cluster with the lowest, statistically significant (FDR <=0.1) cosine distance.

### Identifying druggable gene candidates and prioritizing drugs

Correlation of essentiality is calculated across all 18,133 gene pairs using the rcorr function in the Hmisc package for cell lines that match each of the patient clusters separately. Cell line cluster identity is identified using the previously described methodology. Gene pairs are then filtered so that 1 member of the pair is one of the key TFs in that cluster using TF-score in patient data. We use spearman correlation to lower the frequency of spurious or weak gene essentiality correlations then use a Benjamini-Hochberg adjusted p-value of 0.2 as an initial filter before working with the top 10% of pairs that pass that filter. Pairs are scored individually using the formula below:

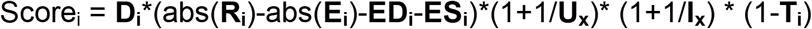

**R_i_** represents the absolute value of the correlation between TF_x_ and Gene_i_.

**D_i_** is a Boolean indicating whether Gene_i_ is targetable by known drugs.

**E_i_** is the median essentiality of Gene_i_ across all cell lines. Essential genes have lower (negative) values.

**ES_i_** is the median essentiality of Gene_i_ in cell lines matching that cluster

**ED_i_** is the differential essentiality of Gene_i_ between cell lines matching that cluster and all cell lines

**U_x_** is a measure of TF_x_ ubiquity across all patient clusters.

**I_x_** is a measure of TF_x_ importance in patient cluster matching cell lines under consideration, this is a ranked variable with higher rank representing less importance.

**T_i_** is a measure of how many drugs that target Genei also affect genes that are broadspectrum essential across all cell lines.

Drug candidates are prioritized using the following formula:

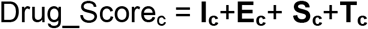

**I_c_** represents the nature of the interaction between compound_c_ and gene_i_. This is a categorical variable converted to an integer with antagonists receiving positive values, agonists and activators receiving negative values.

**E_c_** represents the evidence supporting the interaction between compound_c_ and gene_i_.

This is a categorical variable converted to an integer. Interactions sourced from clinical trials and/or the FDA are given higher ranks (lower values of 1-E_c_) while interactions from single publications and less curated datasets receive lower ranks.

**S_c_** represents the specificity of compound_c_ compared to all other compounds in dataset.

**S_c_** is defined as:

1-(# of genes targeted by compound_c_)/median(# of genes targeted by all compounds in the dataset).

Higher values of Sc are desirable.

**T_c_** represents the median inferred toxicity of compound_c_ based on all predicted targets of compound_c_. The toxicity is inferred from the median gene dependency of all genes targeted by that compound across all cell lines. Lower (more negative) values of Tc imply a more toxic compound.

### Validating drug candidates in vitro

Cells were seeded in 96 well plate at a density of 3000 cells/well. HCC1143 cells were used as a control across all experiments. A498 (ATCC-HTB-44) cells were treated with half log dilutions of RG-7112 (Selleckchem S7030). Cells were assessed using Cell Titer Glo (Promega-G9242) after 6h. DMS53 (ATCC-CRL-2062) cells were treated with log dilutions of Mercaptopurine Hydrate (Cayman Chemicals 23675) and half log dilutions of Teglarinad Chloride (MedChemExpress – HY-12334). Cells were assessed with Cell Titer Glo at 6h and 48h respectively. HEPG2 cells were treated with doubling dilutions of TP-472 (Tocis-6000) and were assessed after 48h on treatment. All assays were performed with 3 biological replicates consisting of 3 technical replicates each at every drug concentration. To minimize plate variation, the control cell line was added to the same 96 well. Internal controls also included wells incubated with vehicle and medium alone.

